# IL-1β and TNF drive endothelial dysfunction and coagulopathy in acute COVID-19

**DOI:** 10.64898/2026.03.21.713333

**Authors:** Helen Mostafavi, Brittany Hill, Christina Nalkurthi, Stefanie M Bader, Yanshan Zhu, Alexander Yu, Philip M. Hansbro, Marcel Doerflinger, Matt D. Johansen, Kirsty R. Short, Keng Yih Chew, Emma J Gordon, Larisa I Labzin

## Abstract

Vascular dysfunction and coagulopathy are hallmarks of severe COVID-19. How SARS-CoV-2 infection drives endothelial dysfunction, despite the virus not infecting or replicating in endothelial cells, remains controversial. Here, we used an *in vitro* co-culture model of the human pulmonary epithelial-endothelial cell barrier to investigate which inflammatory mediators drive endothelial dysfunction during SARS-CoV-2 infection. SARS-CoV-2 infection of primary human bronchial epithelial cells increased adjacent endothelial cell expression of the leukocyte adhesion marker ICAM-1, disrupted endothelial VE-cadherin junctions, promoted endothelial cell death, and promoted platelet adherence to gaps in the endothelial monolayers. Dexamethasone treatment rescued these dysregulated endothelial phenotypes in infected co-cultures, confirming that inflammatory signalling was the primary driver of SARS-CoV-2-induced endothelial dysfunction. Specifically, epithelial-derived TNF and IL-1β promoted endothelial dysfunction, as inhibition of TNF or IL-1R signalling blocked SARS-CoV-2-induced endothelial dysfunction in co-cultures. SARS-CoV-2-infected wild-type mice, but not TNF, IL-1β, or TNF/IL-1β− deficient mice, displayed increased endothelial ICAM-1 expression, while an anti-IL-1β monoclonal antibody prevented SARS-CoV-2-induced ICAM-1 expression and fibrin deposition in aged K18-ACE2 mice. Our data indicate that TNF and IL-1β are the specific cytokines that drive multiple aspects of endothelial dysfunction during acute SARS-CoV-2 infection, and that inhibiting their signalling pathways may provide therapeutic benefit in preventing vascular complications of COVID-19.

## Introduction

Endothelial cells line blood vessel walls and regulate tissue homeostasis, vessel permeability, immune cell extravasation into the tissue, and clot formation.^1^ Endothelial dysfunction and injury are hallmarks of severe COVID-19, the disease caused by infection with Severe Acute Respiratory Syndrome Coronavirus 2 (SARS-CoV-2)^2^. Clinically, endothelial dysfunction in COVID-19 manifests as oedema, endothelitis, disseminated intravascular coagulation, and thromboses^3–5^, leading to respiratory failure and an increased risk of venous thromboembolism^6^. Currently, prophylactic dose anticoagulation therapy is recommended for all patients with critical COVID-19, even though the evidence that this therapy reduces thrombotic complications, length of hospital stay, or all-cause mortality is limited^7^. Whether non-coagulant interventions protect against endothelial dysfunction and injury in COVID-19 is unknown^7^.

The mechanisms underlying how SARS-CoV-2 infection causes endothelial dysfunction are controversial; however, there are two leading hypotheses. The ‘direct’ hypothesis suggests that SARS-CoV-2 directly infects endothelial cells, causing endothelial dysfunction^9^. Early in the COVID-19 pandemic, initial studies suggested that SARS-CoV-2 displayed endothelial tropism *in vivo*, potentiated by ACE2 expression on a wide variety of endothelial cells^4,10,11^. However, in-depth *in vitro* studies previously performed by our group and others showed that endothelial cells express only trace levels of ACE2 and that direct infection with SARS-CoV-2 is neither productive nor causes endothelial dysfunction^12,13^. Gao et al. recently confirmed that endothelial SARS-CoV-2 infection is not the underlying cause of COVID-19-associated endothelial dysfunction in mice, as mice deficient in endothelial ACE2 displayed the same SARS-CoV-2-induced increases in pulmonary thrombi and endothelial inflammatory marker expression as wild-type (WT) mice^14^.

In contrast, the ‘indirect’ hypothesis posits that the inflammatory cytokine milieu triggered by local SARS-CoV-2 infection causes endothelial inflammation and injury^15^. Our previous work supported this alternative hypothesis of an indirect, inflammatory cause of endothelial dysfunction during SARS-CoV-2 infection^12^. Specifically, we showed in an *in vitro* co-culture model that epithelial infection increased expression of intercellular adhesion molecule 1 (ICAM-1) on adjacent endothelial cells^12^, consistent with other *in vitro* studies demonstrating that epithelial infection causes adjacent endothelial inflammation, loss of barrier function, and endothelial cell death^16–18^. However, the specific inflammatory factors driving endothelial dysfunction during acute COVID-19 infection remain unclear.

Excessive cytokine production and signalling drive severe COVID-19 pathogenesis. Multiple cytokines, including the pro-inflammatory triad of tumour necrosis factor (TNF), Interleukin (IL)-6, and IL-1β, are associated with increased COVID-19 disease severity ^21,22,19^. The broad-spectrum anti-inflammatory drug dexamethasone remains the primary recommended treatment for severe COVID-19, with anti-IL-6 receptor (IL-6R) monoclonal antibody (tocilizumab) and the JAK/STAT signalling inhibitor baricitinib, recommended as further anti-inflammatory treatments^23^. While blocking IL-6 signalling shows some protection from vascular leakage and endothelial glycocalyx loss in human cells in vitro^18,19,24^, whether IL-6 can also promote coagulation and endothelitis remains unexplored. Similarly, whether other cytokines beyond IL-6, such as TNF and IL-1β, contribute to SARS-CoV-2-induced vascular dysfunction remains unclear.

Here, we used treated co-cultures of primary human epithelial and endothelial cells with clinically available immunomodulators and used functional readouts of endothelial dysfunction and coagulation to investigate which cytokines drive endothelial dysfunction. We find that TNF and IL-1β are the central orchestrators of endothelial dysfunction in acute COVID-19, both in human primary cells and in murine models of SARS-CoV-2 infection.

## Results

### Primary epithelial cell SARS-CoV-2 infection drives adjacent endothelial cell dysfunction

To interrogate how epithelial SARS-CoV-2 infection can cause endothelial dysfunction, we established co-cultures of differentiated primary human bronchial epithelial cells (NHBE) with primary human lung microvascular endothelial cells (HMVEC-L) (Fig. S1A). After differentiating the NHBE at an air-liquid interface in the apical compartment of transwells, we seeded the basal side of the transwells with HMVEC-L. We confirmed that the NHBE formed a pseudostratified epithelial layer, while the HMVEC-L formed an uninterrupted endothelial monolayer (Fig. S1B). Compared with undifferentiated NHBE, differentiated NHBE had increased expression of the epithelial marker of ciliated cells, *FOXJ1* (Fig. S1C), increased acetylated tubulin staining (a marker of stable cilia), and increased expression of the mucin-producing enzyme MUC5AC (Fig. S1D). Consistent with our previous observations in Calu3/HMVEC-L co-cultures^12^, at 72 hours post-infection, we detected SARS-CoV-2 nucleocapsid protein (NP) in NHBE cells from infected co-cultures, but not in HMVEC-L cells (Fig. S1E, S1F). SARS-CoV-2 replicated in NHBE but not in HMVEC-L, as we detected increased viral titres in the apical compartment of the NHBE co-cultures, but not in the basal HMVEC-L compartment, or in HMVEC-L monocultures (Fig. S1G).

To test if NHBE infection activates adjacent HMVEC-L, we measured key functional markers of endothelial dysfunction and thrombosis induced by COVID-19 patient serum: increased ICAM-1 expression^25^, increased vascular permeability^26^, endothelial cell death^27^, and platelet adherence to the endothelium^25^. The HMVEC-L of infected co-cultures significantly increased ICAM-1 expression by 72h post-infection (Fig. 1A, 1B), consistent with our previous observations in Calu3/HMVEC-L co-cultures^12^ and COVID-19 patient serum^25^. In contrast, HMVEC-L monocultures exposed to SARS-CoV-2 did not increase ICAM-1 at 72h post-infection (Fig. S2A, S2B). To test whether SARS-CoV-2 infection also affected endothelial junctional integrity, we stained for vascular endothelial cadherin (VE-cadherin). Loss of VE-cadherin localisation at cell-cell junctions indicates reduced vascular monolayer integrity and a subsequent increase in vascular permeability^28^. In mock-infected co-cultures, VE-cadherin localised to intercellular junctions, and the HMVEC-L formed a tight monolayer with no visible gaps between cells (Fig. 1C). In contrast, visible gaps appeared in the HMVEC-L monolayers in the infected co-cultures by 48h post-infection and persisted to 72h post-infection (Fig. 1C, 1D). As with ICAM-1, when HMVEC-L were exposed to SARS-CoV-2 in the absence of epithelial cells, we observed no change in VE-cadherin localisation or in the formation of gaps between endothelial cells compared to mock (Fig. S2C, S2D). Together, these data show that epithelial infection with SARS-CoV-2 activates neighbouring endothelial cells, increasing adhesion molecule expression and disrupting VE-cadherin-mediated monolayer integrity.

**Figure 1.**
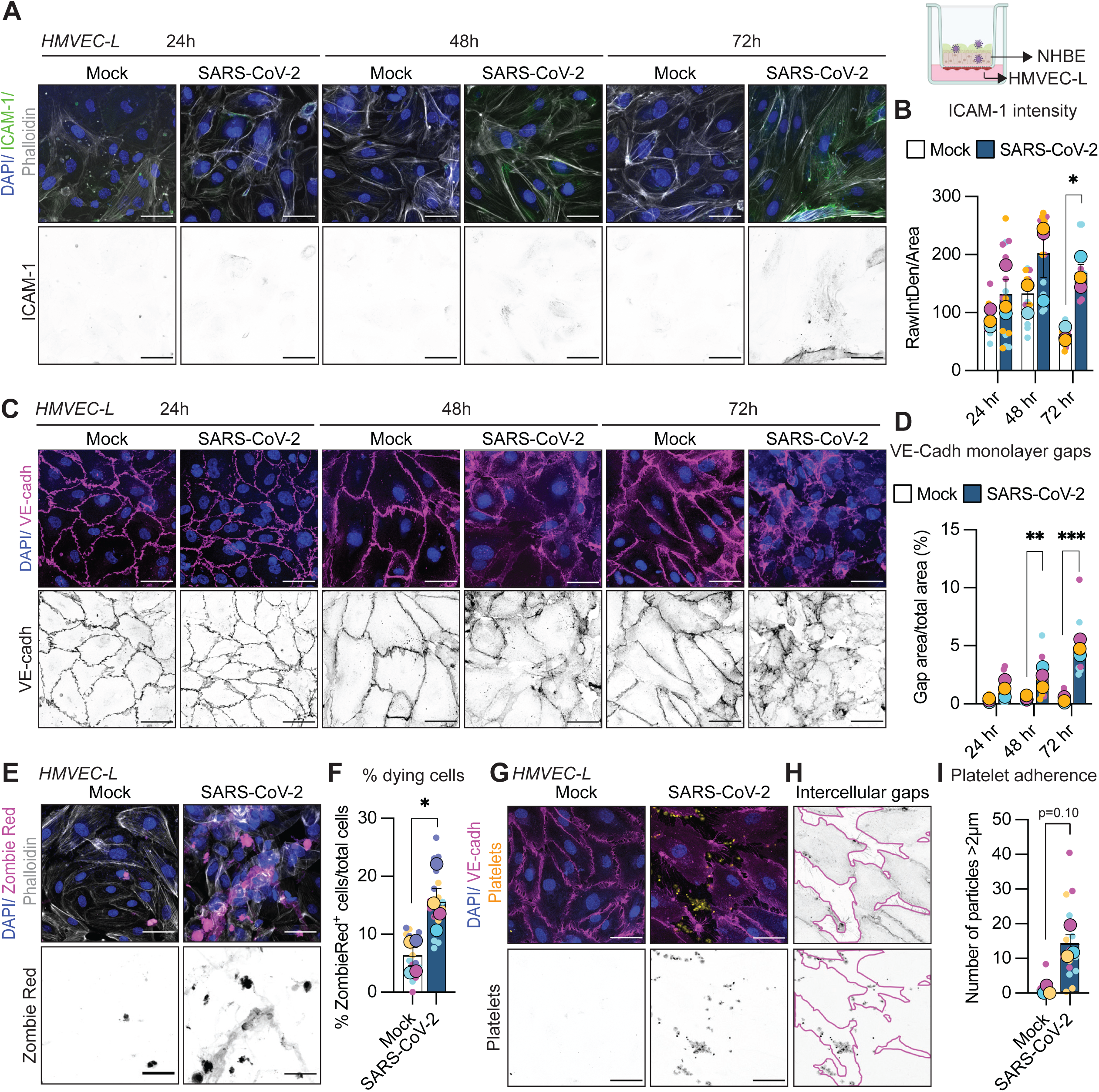
Epithelial infection drives HMVEC-L dysfunction during SARS-CoV-2 infection. **(A-D)** NHBE/HMVEC-L co-cultures were infected with SARS-CoV-2 (MOI 1) or mock-infected, and cells were fixed at 24, 48, and 72h post-infection. **A)** Immunofluorescence staining of ICAM-1 (green), F-actin (phalloidin; grey), and nuclei (DAPI; blue) in HMVEC-L in the basal compartment of the co-culture. Images are representative of three independent experiments. **B)** Quantification of HMVEC-L ICAM-1 intensity analysed by 2-way ANOVA with Sidak’s multiple comparison’s test. **C)** Immunofluorescence staining of VE-cadherin (magenta) in HMVEC-L in the basal compartment of the co-culture. Images are representative of three independent experiments. **D)** Quantification of the percentage of gaps in the endothelial monolayer was analysed by 2-way ANOVA with Sidak’s multiple comparison’s test. **E-H)** NHBE/HMVEC-L co-cultures were infected with SARS-CoV-2 (MOI 1) or mock-infected, and cells were fixed at 72h post-infection. **E)** Immunofluorescence staining of Zombie Red (magenta) in HMVEC-L in the basal compartment of the co-culture. Images are representative of four independent experiments. **F)** The percentage of Zombie Red-positive cells between conditions was analysed using the Kruskal-Wallis test with Dunn’s multiple testing correction. **G)** Immunofluorescence staining and imaging of CellTrace Yellow-labelled platelets (yellow), VE-cadherin (magenta), and nuclei (DAPI; blue) in HMVEC-L in the basal compartment of the co-culture. Images are representative of three independent experiments. **H)** Gaps in the cellular monolayer were outlined relative to VE-cadherin staining, and only the platelets present in gaps were quantified. **I)** The number of platelets (CellTrace Yellow positive particles larger than 2 mm) in gaps was quantified and analysed by the Kruskal-Wallis test with Dunn’s multiple testing correction. Scale bar for all images = 50 µm. 5 ROIs per experiment were quantified (small dots) and are colour-coded per experiment. The average of the 5 ROIs is represented by the large dot (colour-coded by experiment), and the data show the mean ± SEM. Asterisks indicate statistical significance: * *p* < 0.05, ** p < 0.01, *** p < 0.001.

Endothelial cell death is prevalent in severe COVID-19 and may precede thrombosis^27^. We observed significant HMVEC-L cell death in our infected co-cultures at 72h post-infection (Fig. 1E, 1F), even though the HMVEC-L cells were not positive for SARS-CoV-2 infection (Fig. S1E). HMVEC-L without epithelial cells exposed to SARS-CoV-2 did not undergo cell death (Fig. S2E, S2F). We hypothesised that endothelial cell death and loss of intercellular junctional integrity (Fig. 1C, 1E) may promote platelet adherence to the endothelium as part of the haemostatic process to plug gaps in disrupted endothelial monolayers. When we added platelets to the basal compartment of our co-cultures, we observed that platelets accumulated in the SARS-CoV-2-infected co-cultures, but not in the mock-infected co-cultures (Fig. 1G), consistent with Weener et al’s observations that endothelial cells treated with plasma from COVID-19 patients, but not healthy controls, promote platelet aggregation^25^. These platelets primarily adhered to gaps between endothelial cells (Fig. 1H, 1I). No platelets adhered to HMVEC-L exposed to SARS-CoV-2 in the absence of epithelial cells (Fig. S2G, S2H). These data suggest that epithelial infection with SARS-CoV-2 is sufficient to drive adjacent endothelial cell death and trigger platelet adherence to gaps in the endothelial monolayer, potentially as an early step in thrombus formation.

### Blocking inflammation via dexamethasone prevents endothelial dysfunction

Our data indicated that epithelial cells release soluble factors that act on endothelial cells to drive dysfunction. To test if this soluble factor was likely an inflammatory factor, such as a cytokine or an immunostimulatory host or viral molecule, we treated the co-cultures with the broad-spectrum anti-inflammatory drug dexamethasone. Dexamethasone treatment reduced SARS-CoV-2-induced HMVEC-L ICAM-1 expression (Fig. 2A, 2B) and prevented loss of VE-cadherin junctional integrity (Fig. 2C, 2D), indicating that epithelial infection drives endothelial dysfunction through an inflammatory mediator. Consistent with this hypothesis, dexamethasone treatment did not affect viral replication in epithelial cells (Fig. 2E). In the co-culture system, dexamethasone may block epithelial cytokine release, which could, in turn, prevent endothelial cell activation, or it may directly inhibit endothelial cell signalling. To test if dexamethasone directly inhibited endothelial signalling, we collected SARS-CoV-2-infected NHBE basolateral supernatants, added dexamethasone or PBS (mock), and transferred treated supernatants to HMVEC-L monocultures (Fig. 2F). Dexamethasone moderately blocked NHBE supernatant transfer-induced ICAM-1 expression in HMVEC-L (Fig. 2G, 2H) and prevented the loss of VE-cadherin junctional integrity (Fig. 2I, 2J). Taken together, this data indicates that SARS-CoV-2-infected epithelial cells release inflammatory factors that drive endothelial cell dysfunction, and that dexamethasone can directly act on endothelial cells to prevent inflammatory endothelial activation.

**Figure 2:**
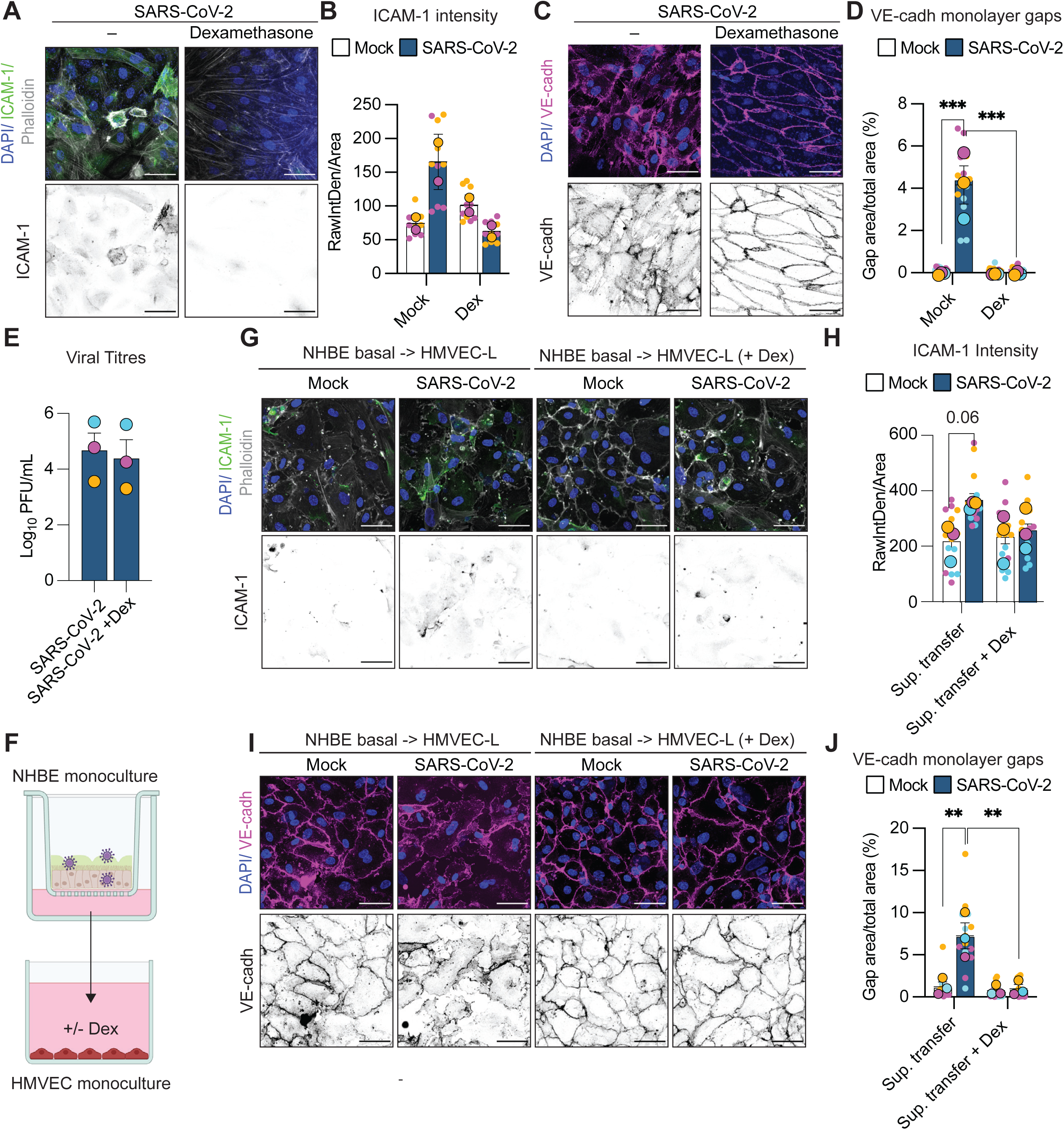
Dexamethasone rescues endothelial function: **(A-E)** NHBE/HMVEC-L co-cultures infected with SARS-CoV-2 (MOI 1) and then treated with 100 mg/mL dexamethasone or media alone immediately post-infection. Cells were fixed at 72h post-infection. **A)** Immunofluorescence staining of ICAM-1 (green) in HMVEC-L in the basal compartment of the co-culture. Images are representative of 2 independent experiments. **B)** Quantification of HMVEC-L ICAM-1 intensity, where data shows mean + SD. **C)** Immunofluorescence staining of VE-cadherin (magenta) in HMVEC-L in the basal compartment of the co-culture. Images are representative of three independent experiments. **D)** Quantification of the percentage of gaps in the endothelial monolayer was analysed by one-way ANOVA with Sidak’s multiple comparison’s test. **E)** Viral titres from the apical compartment of SARS-CoV-2-infected NHBE/HMVEC-L co-cultures, untreated or treated with 100 mg /mL dexamethasone, at 72h post-infection. n = 3 independent experiments, analysed by unpaired, two-way *t*-test. **F)** Schematic of supernatant transfer experiment. **G-J)** NHBE monocultures were infected with SARS-CoV-2 for 48h. The supernatant from the basal compartment was then transferred onto HMVEC-L. 100 mg /mL dexamethasone or PBS was added to the NHBE basal supernatants before they were transferred onto HMVEC-L. After 24h, HMVEC-L were fixed for immunofluorescence staining. **G)** Immunofluorescence staining of ICAM-1 (green) in HMVEC-L. Images are representative of 3 independent experiments. **H)** Quantification of HMVEC-L ICAM-1 intensity analysed by one-way ANOVA with Sidak’s multiple comparison’s test. **I)** Immunofluorescence staining of VE-cadherin (magenta) in HMVEC-L. Images are representative of three independent experiments. **J)** Quantification of the percentage of gaps in the endothelial monolayer was analysed by one-way ANOVA with Sidak’s multiple comparison’s test. Scale bar for all images = 50 µm. 5 ROIs per experiment were quantified (small dots) and are colour-coded per experiment. The average of the 5 ROIs is represented by the large dot (colour-coded by experiment), and the data show the mean ± SEM. Asterisks indicate statistical significance: * *p* < 0.05, ** p < 0.01, *** p < 0.001.

### Epithelial-derived TNF drives endothelial dysfunction

We hypothesised that infected epithelial cells were releasing a cytokine or multiple cytokines that activated endothelial cells. IL-6 can signal in cis through the IL-6 receptor (IL-6R), or in trans by combining with the soluble IL-6R, but only trans IL-6 signalling (i.e., IL-6 + sIL-6R) induces vascular permeability in endothelial cells^19^. TNF is a known driver of endothelial dysfunction^20^, and IL-1β signalling can also trigger vascular permeability *in vitro* and *in vivo*^19,29^. We therefore tested whether IL-6, TNF, or IL-1β signalling was sufficient to phenocopy the endothelial dysfunction we observed after epithelial SARS-CoV-2 infection (Fig. 1). TNF and IL-1β treatment induced robust ICAM-1 expression in HMVEC-L, in contrast to IL-6 (either alone or in combination with sIL-6R) (Fig. S3A, S3B). TNF, IL-1β, and IL-6 + sIL-6R treatment, but not IL-6 alone, disrupted VE-cadherin junctions, consistent with published observations (Fig. S3C, Fig. S3D). TNF and IL-1β also triggered robust endothelial cell death, unlike IL-6 or IL-6 + sIL-6R (Figure S3E, S3F). Finally, while IL-6 + sIL-6R triggered some platelet adherence, TNF and IL-1β treatment both triggered robust platelet adherence, to a similar degree as observed in the infected co-cultures (Fig. S3G, S3H). TNF and IL-1β signalling are therefore sufficient to drive the same endothelial dysfunction phenotypes we observed in SARS-CoV-2 infection. In contrast, IL-6 signalling disrupts VE-Cadherin junctions and platelet adherence but does not drive endothelial ICAM-1 expression or cell death.

To test if TNF was the soluble factor driving endothelial dysfunction in the co-cultures, we therefore blocked TNF signalling with an anti-TNF monoclonal antibody (Adalimumab), which prevents TNF from binding to and signalling through the TNF receptor^30^. Adalimumab completely inhibited SARS-CoV-2-induced ICAM-1 expression (Fig. 3A, B), prevented loss of VE-cadherin junctional integrity (Fig. 3C, D), blocked endothelial cell death (Fig. 3E, F), and prevented platelet adherence to the endothelium (Fig. 3G, H). TNF is therefore a primary driver of endothelial dysfunction in SARS-CoV-2-infected co-cultures.

**Figure 3.**
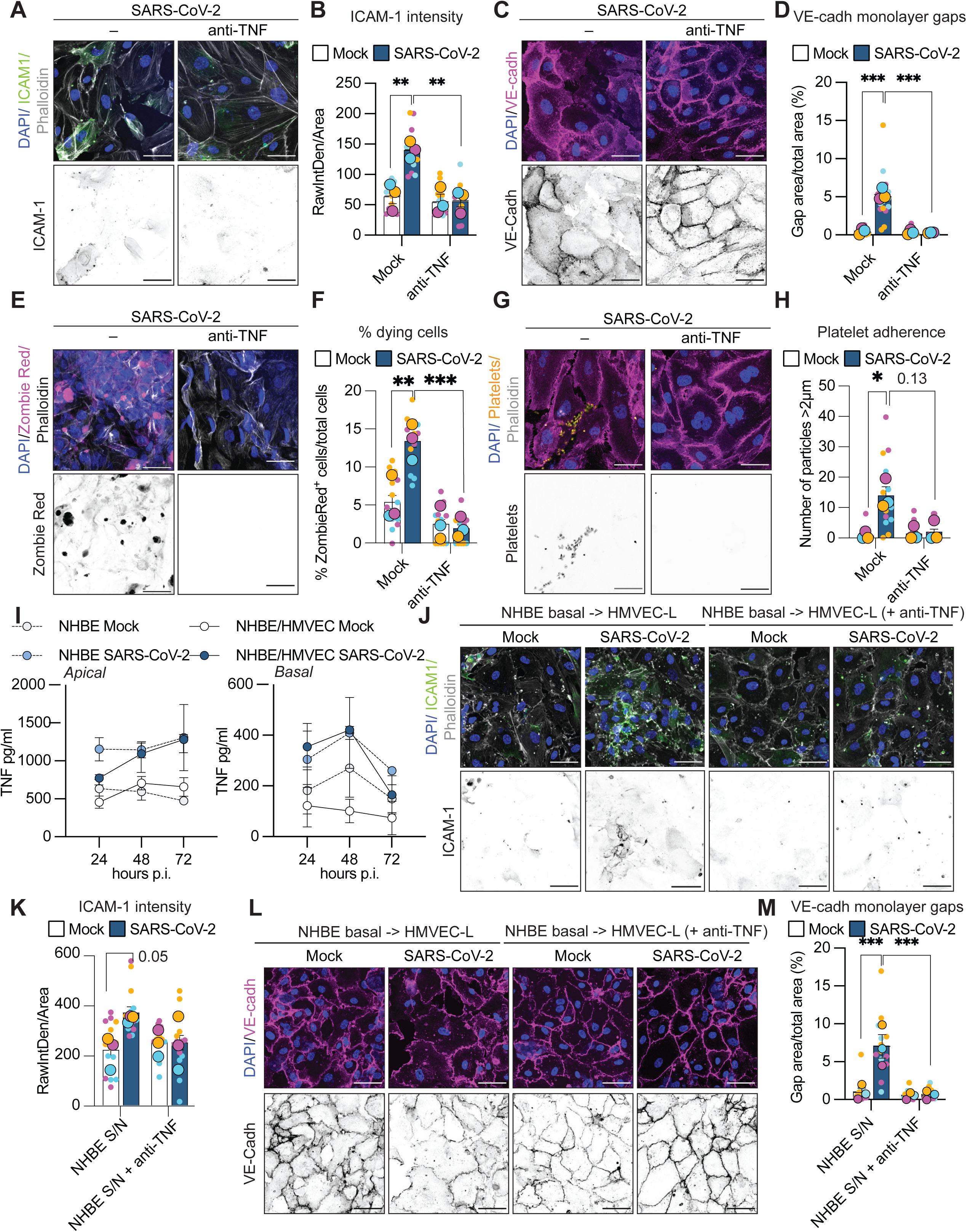
TNF drives HMVEC-L dysfunction in SARS-CoV-2-infected co-cultures. **(A-H)** NHBE/HMVEC-L co-cultures were infected with SARS-CoV-2 (MOI 1) or mock-infected and treated with 10 mg/mL anti-TNF (Adalimumab) immediately post-infection. Cells were fixed at 72h post-infection. **A)** Immunofluorescence staining for ICAM-1 (green), F-actin (Phalloidin; grey), and nuclei (DAPI; blue). Images are representative of n = 3 independent experiments. **B)** ICAM-1 intensity between conditions was analysed by one-way ANOVA, with Sidak’s multiple comparison test. **C)** Immunofluorescence staining for VE-cadherin (magenta), F-actin (Phalloidin; grey), and nuclei (DAPI; blue). Images are representative of n = 3 independent experiments. **D)** Quantification of gaps in the endothelial monolayer under different conditions was determined by calculating the percentage of the image area covered by gaps, and analysed by one-way ANOVA, with Sidak’s multiple comparison’s test. **E)** Zombie Red-stained cells (magenta) indicate cells (containing F-actin and nuclei) undergoing cell death. Images are representative of n = 3 independent experiments. **F)** The percentage of Zombie Red positive cells was analysed by one-way ANOVA, with Sidak’s multiple comparison’s test. **G)** Immunofluorescent staining of CellTrace Yellow-labelled platelets incubated with HMVEC-L. Images are representative of n = 3 independent experiments. **H)** The number of platelets (CellTrace Yellow positive particles larger than 2 mm) in gaps was quantified and analysed by the Kruskal-Wallis test with Dunn’s multiple testing correction. **I)** NHBE monocultures and NHBE/HMVEC-L co-cultures were infected with SARS-CoV-2 (MOI 1) or mock-infected, and TNF levels in the apical and basal supernatants were analysed at 24, 48, and 72h post-infection. Data show the mean ± SEM of 3 independent experiments, analysed by 2-way ANOVA. **(J-M)** NHBE monocultures were infected with SARS-CoV-2 for 48h. The supernatant from the basal compartment was then transferred onto HMVEC-L. Anti-TNF (10 mg/mL) or PBS was added to the NHBE basal supernatants before they were transferred onto HMVEC-L. After 24h, HMVEC-L were fixed for immunofluorescence staining. **J)** Immunofluorescence staining of ICAM-1 (green) in HMVEC-L. Images are representative of 3 independent experiments. **K)** Quantification of HMVEC-L ICAM-1 intensity was analysed by one-way ANOVA with Sidak’s multiple comparison’s test. **L)** Immunofluorescence staining of VE-cadherin (magenta) in HMVEC-L. Images are representative of three independent experiments. **M)** Quantification of the percentage of gaps in the endothelial monolayer was analysed by one-way ANOVA with Sidak’s multiple comparison’s test. Scale bar for all images = 50 µm. 5 ROIs per experiment were quantified (small dots) and are colour-coded per experiment. The average of the 5 ROIs is represented with the large dot (colour-coded per experiment), and the data shows mean ± SEM. Asterisks indicate statistical significance: * *p* < 0.05, ** p < 0.01, *** p < 0.001.

To confirm that the infected epithelial cells are the source of TNF in the co-cultures, we compared TNF levels in NHBE monocultures versus NHBE/HMVEC-L co-cultures. SARS-CoV-2 infection triggered the same magnitude of TNF release from NHBE alone as from NHBE/HMVEC-L co-cultures, across multiple timepoints and in both the apical and basal compartments (Fig. 3I). To test if epithelial-derived TNF was sufficient to drive endothelial dysfunction, we transferred the basal supernatant of infected NHBE monocultures onto HMVEC-L monocultures and added anti-TNF directly to the HMVEC-L. Anti-TNF treatment blocked supernatant-transfer-induced increases in endothelial ICAM-1 (Fig 3J, 3K) and disruptions in VE-cadherin junctions (Fig 3L, 3M). These supernatant transfer results phenocopied our co-culture experiments. Finally, we confirmed that 400pg/ml TNF, which was approximately the level of TNF we detected in the basal compartment of infected co-cultures at 48h post infection (Fig 3I), was sufficient to induce endothelial ICAM1 expression (Fig. S4A, S4B), loss of VE-cadherin junctions (Fig. S4C, S4D), endothelial cell death (Fig. S4E, S4F) and platelet adherence (Figure S4G, S4I). Together, these experiments confirm that epithelial-derived TNF is sufficient to drive endothelial dysfunction during SARS-CoV-2 infection.

### Inhibiting IL-1 signalling with Anakinra prevents endothelial dysfunction

IL-1β signalling was also sufficient to drive the full gamut of endothelial dysfunction phenotypes we observed in the co-culture experiments (Supp. Fig. 3). We therefore tested whether blocking IL-1R signalling with the recombinant IL-1R antagonist Anakinra^31^ would affect SARS-CoV-2-induced endothelial dysfunction in our co-culture system. Anakinra treatment of the co-cultures completely blocked SARS-CoV-2-induced endothelial ICAM-1 expression (Fig. 4A, B), prevented loss of VE-cadherin junctional integrity (Fig. 4C, D), and inhibited endothelial cell death (Fig. 4E, F), phenocopying our findings with anti-TNF treatment (Fig. 3A-H). This data suggests that IL-1 signalling also drives SARS-CoV-2-induced endothelial dysfunction in the epithelial endothelial co-cultures.

**Figure 4.**
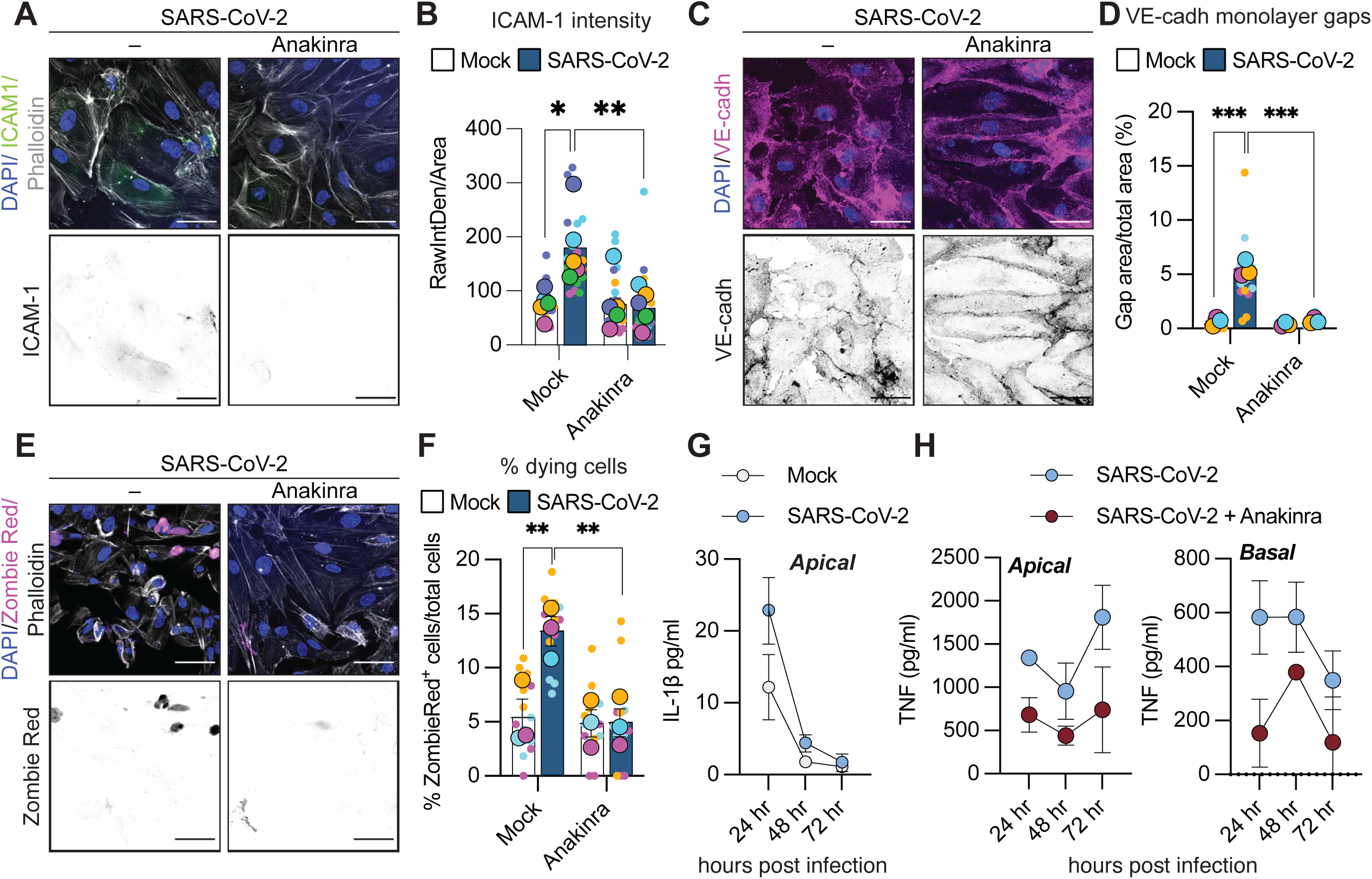
Anakinra also rescues endothelial function in SARS-CoV-2-infected co-cultures. **(A-F)** NHBE/HMVEC-L co-cultures were infected with SARS-CoV-2 (MOI 1) or mock-infected and treated with 10 mg/mL Anakinra immediately post-infection. Cells were fixed at 72h post-infection. **A)** Immunofluorescence staining for ICAM-1 (green), F-actin (Phalloidin; grey), and nuclei (DAPI; blue). Images are representative of n = 5 independent experiments. **B)** ICAM-1 intensity between conditions was analysed by one-way ANOVA, with Sidak’s multiple comparison test. **C)** Immunofluorescence staining for VE-cadherin (magenta), F-actin (Phalloidin; grey), and nuclei (DAPI; blue). Images are representative of n = 3 independent experiments. **D)** Quantification of gaps in the endothelial monolayer under different conditions was determined by calculating the percentage of the image area covered by gaps, and analysed by one-way ANOVA, with Sidak’s multiple comparison’s test. **E)** Zombie Red-stained cells (magenta) indicate cells (containing F-actin and nuclei) undergoing cell death. Images are representative of n = 3 independent experiments. **F)** The percentage of Zombie Red positive cells was analysed by one-way ANOVA, with Sidak’s multiple comparison’s test. **G)** IL-1β levels in the apical supernatant of NHBE monocultures infected with SARS-CoV-2 (MOI 1) or mock-infected, at 24, 48, and 72h post-infection (n = 3 independent experiments). **H)** TNF levels in the apical (left panel) and basal (right panel) supernatant of SARS-CoV-2-infected NHBE/HMVEC-L co-cultures, untreated or treated with Anakinra, at 24, 48, and 72h post-infection (n = 2 independent experiments). Scale bar for all images = 50 µm. 5 ROIs per experiment were quantified (small dots) and are colour-coded per experiment. The average of the 5 ROIs is represented with the large dot (colour-coded per experiment), and the data shows mean ± SEM. Asterisks indicate statistical significance: * *p* < 0.05, ** p < 0.01, *** p < 0.001.

Given that IL-1β signalling via IL-1R can induce TNF production^32^, and epithelial-derived TNF was sufficient to drive endothelial dysfunction, we hypothesised that IL-1β may be necessary for SARS-CoV-2-induced TNF production in epithelial cells. We detected a modest increase in apical IL-1β levels in NHBE monocultures at 24h post-infection, which decreased to almost undetectable levels by 48 and 72h post-infection (Fig. 4G). To test whether IL-1 signalling was upstream of TNF release in SARS-CoV-2-infected NHBE cells, we treated the cells with Anakinra and measured TNF release. Anakinra treatment reduced TNF levels in SARS-CoV-2-infected epithelial cells at all time points (Fig. 4H), suggesting that IL-1R signalling may be upstream of TNF signalling. Our data therefore suggest that both IL-1R and TNF signalling drive SARS-CoV-2-induced endothelial dysfunction.

### TNF and IL-1β drive SARS-CoV-2-induced endothelial dysfunction in vivo

While our data suggest that epithelial-derived IL-1β and TNF are sufficient to drive endothelial dysfunction *in vitro*, it is unclear whether they also drive vascular dysfunction *in vivo* during COVID-19. We therefore assessed endothelial dysfunction using two separate murine models. In WT mice infected with the mouse-adapted P21 strain of SARS-CoV-2^33,34^, we observed significant upregulation of ICAM-1 expression in CD31-positive endothelial cells of large pulmonary blood vessels, at the peak of disease on day 3 post-infection (Fig. 5A, B). However, SARS-CoV-2-infected *Tnf^-/-^, Il1b^-/-^* or *Tnf^-/-^/Il1b^-/-^*mice did not increase endothelial ICAM-1 expression in the large blood vessels (Fig. 5A, 5B). Similarly, aged K18-ACE2 mice^35^ infected with SARS-CoV-2 showed significantly increased endothelial ICAM-1 expression in large pulmonary blood vessels on day 6 post-infection (Fig. 5C, D). Inhibiting IL-1β signalling prophylactically or therapeutically with a murine anti-IL-1β antibody prevented SARS-CoV-2-induced ICAM-1 expression (Fig. 5C, 5D).

**Figure 5.**
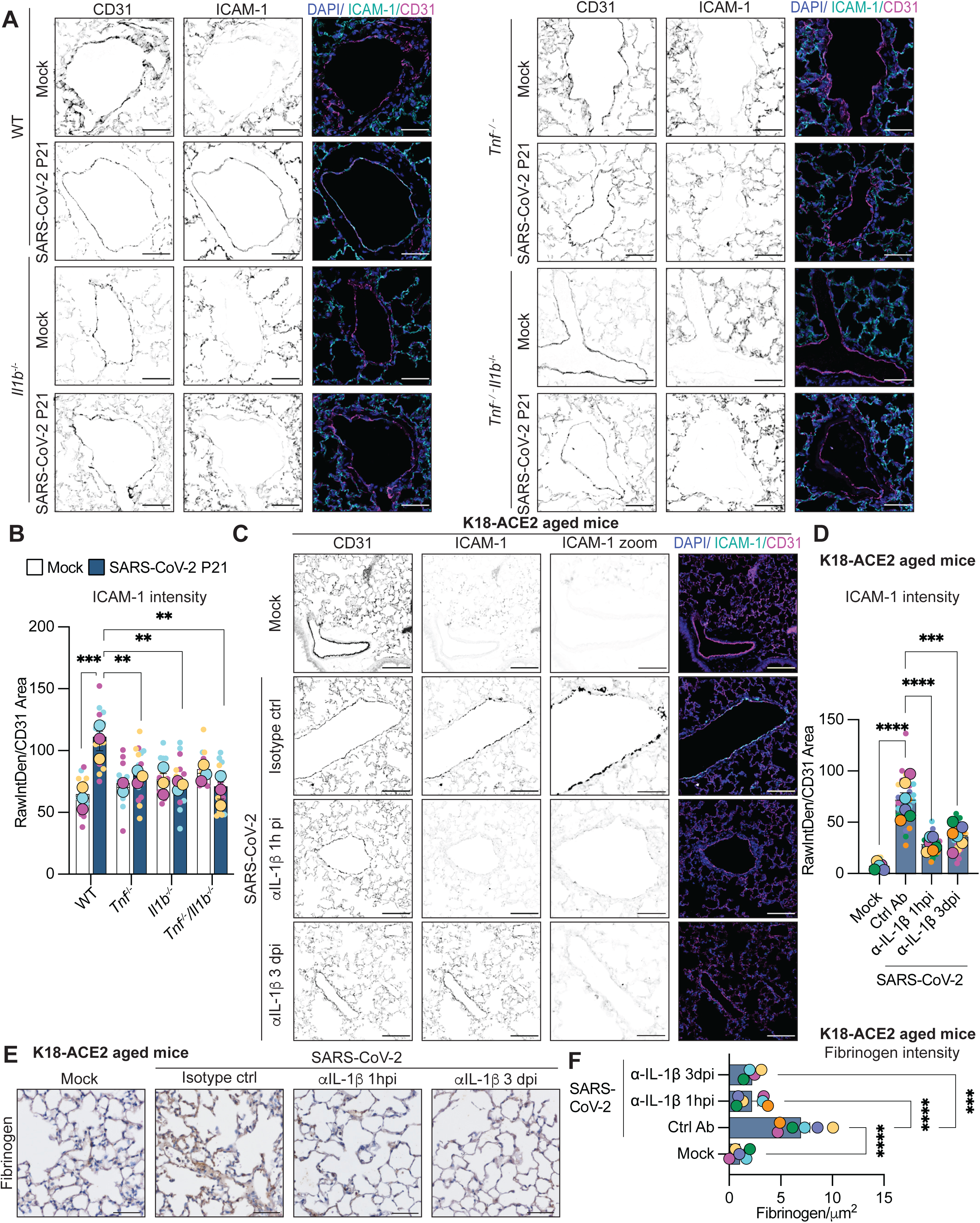
TNF and IL-1β drive SARS-CoV-2 endothelial dysfunction *in vivo*. **A)** Representative immunohistochemistry (IHC) images of lungs from SARS-CoV-2-infected (10^4^ TCID50) or mock-infected wild-type (WT), *Tnf*^-/-^, *Il1b*^-/-^, and *Tnf*^-/-^*IL1b*^-/-^ mice, harvested at 3 days post-infection (dpi). Tissues were stained with CD31 (magenta), ICAM-1 (green), and DAPI (blue). Scale bar = 50 µm. Images are representative of 5 images per mouse, 3 mice per group. **B)** Quantification of ICAM-1 intensity in areas of CD31 staining, analysed by 2-way ANOVA with Sidak’s multiple testing correction. 5 ROIs per mouse were quantified (small dots) and are colour-coded per mouse. The average of the 5 ROIs is represented with the large dot (colour-coded per mouse), and the data shows mean ± SEM. **C-F)** Aged K18-hACE c57BL/6□J mice infected with SARS-CoV-2 (10^3^ PFU) and treated with an isotype control or anti-IL-1β antibody at 1h or 3 days post-infection. Lungs were harvested at day 6 post-infection. **C)** Lung tissues were stained with CD31 (magenta), ICAM-1 (green), and DAPI (blue). Scale bar = 100 µm. Images are representative of 5 images per mouse, 4-6 mice per group. **D)** Quantification of ICAM-1 intensity in areas of CD31 staining, analysed by one-way ANOVA, with Sidak’s multiple comparison’s test. 5 ROIs per mouse were quantified (small dots) and are colour-coded per mouse. The average of the 5 ROIs is represented by the large dot (colour-coded per mouse), and the data show the mean ± SEM. **E)** Lung tissues were stained for fibrinogen. Scale bar = 50 µm. Images are representative of 4-6 mice per group. **F)** Quantification of fibrinogen staining intensity analysed by one-way ANOVA, with Sidak’s multiple comparison’s test. Asterisks indicate statistical significance: * *p* < 0.05, ** p < 0.01, *** p < 0.001.

Excessive fibrin deposition is another hallmark of severe COVID-19 coagulopathy^36^. Fibrin deposition occurs when thrombin cleaves circulating fibrinogen into bioactive fibrin. As fibrin is an integral structural component of clots alongside red blood cells and platelets, its deposition indicates excessive coagulation. SARS-CoV-2-infected aged K18-ACE2 mice displayed significantly increased alveolar fibrin/fibrinogen staining and deposition compared to uninfected mice (Fig. 5E, F). As with ICAM-1, inhibiting IL-1β signalling prevented SARS-CoV-2-induced fibrinogen deposition (Fig. 5E, F). Our data therefore show that IL-1β and TNF signalling are necessary for SARS-CoV-2–induced endothelial activation, and that IL-1β signalling is necessary for SARS-CoV-2-induced fibrin deposition *in vivo*.

## Discussion

Here, for the first time, we demonstrate that TNF and IL-1β drive endothelial adhesion marker expression, vascular permeability, and cell death, and promote thrombosis in acute SARS-CoV-2 infection. Our data suggest that SARS-CoV-2-infected epithelial cells produce IL-1β and TNF. These epithelial-derived cytokines subsequently act on the adjacent endothelium to increase ICAM-1 expression, disrupt VE-cadherin junctions, trigger endothelial cell death, and promote platelet adherence. If the two cytokines acted synergistically to trigger endothelial dysfunction, we would expect only a partial reduction with individual cytokine antagonists and complete reversal of dysfunction with combined anti-TNF and anti-IL-1R treatment. However, given that blocking TNF or IL-1 signalling individually completely reversed SARS-CoV-2-induced endothelial dysfunction, we hypothesise that IL-1β is upstream of TNF within the same signalling pathway. Our *in vitro* results are consistent with our *in vivo* findings in SARS-CoV-2-infected mice, where TNF-and IL-1β-deficient mice did not increase endothelial ICAM-1 expression post-infection, which could underpin our previous findings that *Tnf*^-/-^ and *Il1b^-/-^* animals have improved overall disease outcome^33,34^. Similarly, treatment with an anti-IL-1β antibody decreased SARS-CoV-2-induced endothelial ICAM-1 expression and fibrinogen deposition *in vivo*, in agreement with previous studies showing that Anakinra prevented SARS-CoV-2-induced vascular leak^29^. Our findings provide clear evidence supporting the inflammatory hypothesis of vascular dysfunction in acute COVID-19.

TNF has a well-described role in driving endothelial dysfunction in vascular diseases^20^. Accordingly, TNF blockers currently licensed for clinical use (e.g., Adalimumab, Infliximab) may reduce endothelial complications associated with SARS-CoV-2 infection. Indeed, a study of 600 patients with rheumatic disease and COVID-19 found a significant inverse association between anti-TNF therapy and hospitalisation, even after controlling for confounding factors (OR=0.40, 95% CI 0.19 to 0.81; p=0.01)^37^. However, whether this is directly related to endothelial dysfunction or to coagulation remains unclear. In this regard, there is stronger clinical evidence of dexamethasone’s effectiveness, and we also found that it reduced endothelial dysfunction. Dexamethasone reduces the relative risk of 28-day mortality by approximately 30% in patients with severe COVID-19 requiring mechanical ventilation^38^. More importantly in this context, dexamethasone also reduces biomarkers related to endothelial cell injury (e.g., angiopoietin-2 [Ang-2] and ICAM-1) in severe COVID-19 patients^39^. Furthermore, COVID-19 patients treated with dexamethasone have significantly lower D-dimer levels (a well-established coagulation biomarker) than untreated patients^40^. The efficacy of dexamethasone in treating patients with severe COVID-19 may therefore, at least in part, relate to its ability to restore endothelial cell homeostasis.

In the present study, Anakinra treatment reduced endothelial dysfunction and thrombosis, consistent with previous observations that IL-1β reduced vascular permeability in COVID-19^29^. However, the role of Anakinra in the treatment of COVID-19 remains controversial. Fanlo and colleagues found that Anakinra treatment was not more effective than the standard of care in reducing the need for mechanical ventilation among patients with severe COVID-19 pneumonia^41^. In contrast, Cavalli and colleagues found that in patients with COVID-19 and acute respiratory distress syndrome managed with non-invasive ventilation outside of the ICU, high-dose Anakinra was associated with clinical improvement^42^. Whether these study-to-study differences reflect the differing disease severity between patient cohorts remains to be determined. However, it is interesting to note that in a meta-analysis of 2032 COVID-19 patients, Anakinra treatment was associated with a significant reduction in D-dimer levels^43^. These data underscore the need for future therapeutic studies to measure endothelial dysfunction and coagulation biomarkers to fully assess the treatment’s efficacy.

Our data indicating that both IL-1β and TNF are increased in SARS-CoV-2-infected *in vitro* co-cultures is consistent with published murine models of severe SARS-CoV-2 infection, where both IL-1β and TNF are significantly increased in infected lungs by 3 days post-infection^33^. TNF-deficient, IL-1β-deficient, and IL-1R-deficient mice are all protected from severe SARS-CoV-2 infection^33,34^, indicating that TNF and IL-1 signalling are both necessary to drive severe COVID-19. IL-1R- and IL-1β-deficient mice had significantly lower TNF levels in lung homogenates than WT mice^34^. In contrast, the TNF-deficient mice had equivalent IL-1β levels^33^, supporting our findings that IL-1β is upstream of TNF.

Intriguingly, we detected epithelial cell IL-1β release during SARS-CoV-2 infection, in contrast to Barnett et al. and Planes et al. ^44,45^. This discrepancy may reflect differences in experimental setup, such as timepoints and viral dose or strain, or differences in the sensitivity of the assays used to detect IL-1β. While IL-1β drives SARS-CoV-2-induced disease in mice, many canonical inflammasome components required for IL-1β processing appear dispensable for COVID-19 pathogenesis. Bader et al demonstrated that mice lacking NLR family pyrin domain-containing 3 (NLRP3), NOD-like-receptor containing a Pyrin domain 1 (NLRP1), apoptosis-associated speck-like protein containing a CARD (ASC), Caspase-1, or the pyroptotic effectors Gasdermin D and E, all had the same level of pathology and survival as WT mice during SARS-CoV-2 infection^34^. Therefore, how SARS-CoV-2 infection triggers epithelial IL-1β release remains to be elucidated. We speculate that this epithelial-derived IL-1β subsequently signals in a paracrine manner to trigger epithelial TNF release, sufficient to drive endothelial dysfunction. Airway epithelial cells may therefore be an underappreciated source of TNF and IL-1β during infection *in vivo*. However, our findings do not exclude the possibility that other cells, including immune cells such as macrophages, neutrophils, and T cells, may contribute to the circulating IL-1β and TNF driving endothelial dysfunction during SARS-CoV-2 infection *in vivo*.

During COVID-19, platelet-fibrin clots or haemolysed red blood cells can occlude blood vessels^27^. Apoptotic endothelial cells become pro-coagulant and promote platelet deposition^46^, while haemolysed red blood cells appear to adhere to and ‘plug’ the gaps created by necroptotic endothelial cells^27^. We observed platelets adhering to the gaps in the endothelial monolayer in our SARS-CoV-2-infected transwells. We anticipate that platelets bind to the exposed collagen on the transwell membranes, as occurs *in vivo* when platelets bind to collagen in the basement membrane. Our study showed that blocking TNF and IL-1R signalling prevents endothelial cell death, platelet adhesion, and, in the case of IL-1β, fibrinogen deposition. We therefore predict that TNF and IL-1β play a role in early clot formation by promoting endothelial cell death. As TNF can induce both apoptotic and necroptotic cell death^47^, blocking TNF signalling may provide a therapeutic avenue for preventing platelet-fibrin clots and haemolysed red blood cell occlusions. Finally, while TNF and IL-1β are associated with severe disease in other respiratory infections^48^, the exacerbated coagulation phenotype may be unique to SARS-CoV-2 infection, as the SARS-CoV-2 Spike protein can bind to fibrin, stabilise clot formation, and promote further thromboinflammation by activating complement^49^. Our findings that IL-1β and TNF drive endothelial dysfunction during SARS-CoV-2 infection warrant further studies into the roles of these cytokines in other infections that trigger vascular complications.

The present study was subject to several limitations. Firstly, endothelial cells are a heterogeneous cell population. Accordingly, herein we focused on the role of human lung microvascular endothelial cells in SARS-CoV-2 pathogenesis, using our *in vitro* model to recapitulate the pulmonary epithelial-endothelial barrier, and our *in vivo* studies on the mouse lung. Endothelial cells in other extra-respiratory organs may play a different role in COVID-19, which warrants further investigation.

Secondly, increasing endothelial cell adhesion molecule expression and vascular permeability are important and necessary host defence responses to infection. The increase in ICAM-1 expression and loss of VE-cadherin junctional integrity we observed are consistent with prolonged, dysfunctional responses, rather than protective immune responses^50^. At lower viral doses, we anticipate that epithelial cells will release less IL-1b and TNF; thus, endothelial cells will only transiently express ICAM-1 or open endothelial cell-cell junctions. Indeed, low doses of TNF only transiently promote vascular permeability, while high doses promote permanent vascular leakiness^51^. Identifying the threshold of protective TNF and IL-1 signalling in infection will enable future studies to optimise dosing and timing of immunomodulatory drugs such as Anakinra, Adalimumab, or dexamethasone to prevent vascular damage without compromising host defence.

Finally, the *in vitro* and *in vivo* models used herein did not account for the effect that prior immunity has on SARS-CoV-2 pathogenesis. Accordingly, how vaccination or prior infection affects endothelial function during COVID-19 should be elucidated in future studies.

Nevertheless, the data we present here highlight the important role of endothelial cells and lung epithelial-endothelial cell cross-talk in SARS-CoV-2 pathogenesis. Our data suggest that targeting TNF or IL-1β signalling may prevent vascular dysfunction and clotting in acute COVID-19. Given that IL-1β and TNF are elevated in the serum of long COVID patients^52^, we anticipate that targeting these cytokines may present a novel strategy for alleviating the vascular complications and coagulopathy observed in long COVID^53^.

## Materials and Methods

### Ethics statement

Whole blood from healthy individuals was provided by QML Pathology with written and oral informed consent, and approved by the University of Queensland’s Human Research Ethics Committee (2022_HE002162). All procedures and mouse strains were reviewed and approved by the Walter and Eliza Hall Institute of Medical Research (WEHI) Animal Ethics Committee. They were conducted in accordance with the Prevention of Cruelty to Animals Act (1986) and the Australian National Health and Medical Research Council Code of Practice for the Care and Use of Animals for Scientific Purposes (1997). Studies involving K18-hACE2 aged mice were approved by the Sydney Local Health District (SLHD) Animal Welfare Committee under AWC2020-019, and Institutional Biosafety Committee (IBC) approval was granted under IBC20-051.

### Mice

Male or female wild-type (WT) and gene-targeted mice were bred and maintained in the Specific Pathogen Free (SPF) Physical Containment Level 2 (PC2) Bioresources Facility at WEHI. All mice were on a C57BL/6J background. All procedures involving animals and live SARS-CoV-2 strains were conducted in an OGTR-approved Physical Containment Level 3 (PC3) facility at WEHI (Cert-3621). Mice were transferred to the PC3 laboratory at least 4 days prior to SARS-CoV-2 infection experiments. Animals were age- and sex-matched within experiments (both sexes were used). Experimental mice were housed in individually ventilated microisolator cages under level 3 biological containment conditions with a 12-hour light/dark cycle and provided standard rodent chow and sterile acidified water ad libitum.

Heterozygous male K18-hACE C57BL/6 J mice (strain: 2B6.Cg-Tg(K18ACE2)2Prlmn/J /J) were originally obtained from the Jackson Laboratory. Mice were bred in-house at the Centenary Institute under SPF conditions. Mice were aged to 100 weeks prior to transfer to the Centenary Institute PC3 facility and infection with 10^3^ PFU per mouse (Ancestral; VC01/2020) via intranasal infection. Mice were treated in groups and received either an isotype control antibody (Armenian Hamster IgG Isotype Control; LEI-I-140-100mg; Millenium Science) or anti-mouse IL-1b (LEI-I-437-100mg, Millenium Science). Each mouse received 20 µg antibody via intraperitoneal administration in a 100 µL volume (with PBS). Treatments commenced from 1 hour or 3 days post-infection, and continued daily until the termination of the experiment at day 6 post-infection.

### SARS-CoV-2 strains

For mouse experiments involving mouse-adapted SARS-CoV-2, SARS-CoV-2 VIC2089 clinical isolate (hCoV-19/Australia/VIC2089/2020) was obtained from the Victorian Infectious Disease Reference Laboratory (VIDRL). Viral passages were achieved by serial passage of VIC2089 through successive cohorts of young C57BL/6J (WT) mice. Briefly, mice were infected intranasally with a SARS-CoV-2 clinical isolate. At three days post-infection, mice were euthanised and lungs harvested and homogenised in a Bullet Blender (Next Advance Inc) in 1 mL Dulbecco’s modified Eagle’s medium (DMEM) (Gibco/ThermoFisher) containing steel homogenisation beads (Next Advance Inc). Samples were clarified by centrifugation at 10,000 x g for 5 min before intranasal delivery of 30 µL lung homogenate into a new cohort of naïve C57BL/6 mice. This process was repeated a further 20 times to obtain the SARS-CoV-2 VIC2089 P21 isolate. Lung homogenates from all passages were stored at −80°C.

For experiments involving K18-hACE2 mice, SARS-CoV-2 (Ancestral; VC01/2020) was supplied by the Victorian Infectious Disease Reference Laboratory (VIDLR). SARS-CoV-2 viral titres were generated from clarified supernatant from infected VeroE6 cell cultures and quantified using plaque assays as previously described^54^.

For cell experiments, SARS□CoV□2 isolate hCoV□19/Australia/QLD02/2020 was provided by Queensland Health Forensic & Scientific Services, Queensland Department of Health. Virus was grown on Vero cells and titred as described previously^12^.

### Infection of mice with SARS-CoV-2 P21

Mice were anesthetised with methoxyflurane and inoculated intranasally with 30 μL of DMEM containing SARS-CoV-2. Virus stocks were diluted in serum-free DMEM to a final concentration of 10^4^ TCID50/mouse. After infection, animals were visually checked and weighed daily for at least 10 days. Mice were euthanised at 3 days post-infection by CO_2_ asphyxiation. For histological analysis, animals were euthanised by cervical dislocation. Lungs were collected and stored at −80°C in serum-free DMEM until further processing.

### K18-hACE2 SARS-CoV-2 infection procedures

Mice were lightly anaesthetised with isoflurane and then intranasally inoculated with 30 μL of SARS-CoV-2 (Ancestral; VC01/2020) diluted in endotoxin-free PBS to administer 10^3^ PFU per mouse. Following infection, mice were weighed and clinically scored daily until the completion of the experiment. At day 6 post-infection, mice were euthanised using an intraperitoneal injection of sodium pentobarbitone. The lung tissues were perfused with 0.9% saline solution and then inflated with 10% neutral buffered formalin prior to drop fixing in formalin to inactivate SARS-CoV-2 for 24 hours prior to removal from the PC3 facility.

### NHBE cell culture

Normal human bronchial cells (NHBEs) (Lonza; CC-2540S) guaranteed for air-liquid interface (ALI) culture were expanded and passaged in B-ALI-Growth Basal medium (Lonza; 193516) supplemented with BEGM SingleQuots (Lonza; CC-4175) until passage two at 37 °C, 5% CO_2_. Three NHBE donors were used throughout and were screened to exclude pre-existing co-morbidities (diabetes, heart disease, hypertension, history of smoking). After initial expansion, approximately 50,000 NHBEs (passage 3) were seeded onto 24□well transwell inserts (6.5□mm transwell, 0.4□µm polycarbonate membrane, Corning; COR3470) pre-coated with 0.03 mg/mL Rat Tail Collagen I (Corning; 354236) in the apical and basal compartments. Cells were cultured until a confluent monolayer was observed (approximately 2 days), then airlifted by removing growth medium from the apical compartment. Basal compartment medium was replaced with Pneumacult ALI medium (STEMCELL Technologies; 05001). NHBEs were differentiated for 4-6 weeks with basal media replaced every 2-3 days, until beating cilia and mucus were observed. Fully-differentiated NHBE cultures were subsequently used in co-culture and infection experiments. Details of the NHBE donors used throughout these experiments are listed in Supplementary Table 1.

### HMVEC-L cell culture

Human lung microvascular endothelial cells (HMVEC-L) (Lonza; CC-2527) were cultured in EGM-2MV medium (EBM2 medium supplemented with microvascular endothelial cell growth medium SingleQuots (Lonza; CC-3156 and CC-4147 respectively) until passage two at 37 °C, 5% CO_2_. Five HMVEC-L donors were used throughout and were screened to exclude pre-existing co-morbidities (diabetes, heart disease, hypertension, history of smoking). Monocultures of HMVEC□L were performed on 24□well cell culture transwell inserts (6.5□mm transwell, 0.4□µm polycarbonate membrane, Corning; COR3470) coated with 0.03 mg/mL Rat Tail Collagen I (Corning; 354236). Approximately 50,000 HMVEC□L were seeded per transwell on the apical side and cultured overnight in EGM□2MV medium. Once confluent, HMVEC-L monocultures were cultured for 24h in a 50:50 ratio of ALI-medium and EGM-2MV medium, and then cultured for another 24h in a 50:50 ratio of hydrocortisone-free ALI-medium and EGM-2MV medium before proceeding with the experiment. Details of the HMVEC-L donors used throughout these experiments are listed in Supplementary Table 1.

### NHBE/HMVEC-L co-cultures

NHBE/HMVEC co-cultures were generated using 4-6-week-differentiated NHBE monocultures seeded onto 24-well transwell inserts. Approximately 50,000 HMVEC-L (P3) were seeded onto the basal side of an inverted NHBE transwell and cultured for 1h to allow HMVEC-L to attach to the membrane. Transwells were re-inverted to the normal position and cultured for 24h in 50:50 ALI/EGM-2MV medium. Before infections, co-cultures were cultured for a further 24h in hydrocortisone-free 50:50 ALI/EGM-2MV medium.

### *In vitro* infection with SARS-CoV-2

Cells were infected or mock-infected by adding 100 μL of QLD02 diluted to MOI 1 in hydrocortisone-free 50:50 ALI/EGM-2MV medium or PBS (respectively) into the apical compartment and incubating for 1h at 37°C, 5% CO_2_. The viral inoculum was removed, and PBS was added to the apical and basal compartments (100 μL and 500 μL, respectively) to wash cells. PBS was removed, and basal media was replaced with hydrocortisone-free 50:50 ALI/EGM-2MV medium, supplemented with anti-TNF, Anakinra, or dexamethasone where indicated. Basal media was refreshed every 24h, and cells were incubated at 37°C with 5% CO2. All studies with SARS□CoV□2 were performed under physical containment 3 (PC3) conditions and were approved by The University of Queensland Biosafety Committee (IBC/374B/SCMB/2020).

### Cell stimulations and *in vitro* treatments

HMVEC-L monocultures were stimulated with indicated concentrations of recombinant human TNF (Thermo Fisher Scientific; PHC3011), recombinant human IL-1β (RnD systems; 201-LB), recombinant IL-6 (Peprotech; 200-06-20UG) or recombinant human soluble IL-6R (Peprotech; 200-06RC-20UG) diluted in hydrocortisone-free 50:50 ALI/EGM-2MV medium for 24h at 37 °C, 5% CO_2_.

NHBE monocultures or NHBE/HMVEC-L co-cultures were treated with 10 mg/mL recombinant human monoclonal anti-TNF (Adalimumab, Thermo Fisher Scientific; LT100-1MG), 10 mg /mL recombinant human IL-1 receptor antagonist (Anakinra, Swedish Orphan Biovitrum Pty Ltd), or 100 mg /mL dexamethasone (Sigma-Aldrich; D2915). All treatments were diluted in hydrocortisone-free 50:50 ALI/EGM-2MV medium and added into the basal compartment immediately after removing the viral inoculum (0h post-infection). The medium in the basal compartment was replaced every 24h with media containing freshly diluted treatments throughout the experiments.

### Supernatant transfers

HMVEC-L (passage 3) were cultured in EGM-2MV media. 12-well chamber slides (Ibidi; 81201) were pre-coated with 0.03 mg/mL Rat Tail Collagen I (Corning; 354236) and approximately 50,000 HMVEC-L were seeded per well in EGM-2MV medium. Cells were cultured for 24h at 37°C, 5% CO_2_ to reach confluency, then the medium was changed to 50:50 ALI/EGM-2MV, and cells were cultured for a further 24h. The medium was then changed again to hydrocortisone-free 50:50 ALI/EGM-2MV medium and cell culture for another 24h before proceeding to supernatant transfers. At 48h post-infection, the medium was removed from cells, and supernatant (200 μL) from the basal compartment of mock or infected NHBE monocultures was transferred into the chamber slide wells. Cells were incubated at 37°C in 5% CO_2,_ and the supernatant was harvested after 24h. Cells were then fixed in 4% PFA for 30 min before immunofluorescence staining.

### Immunofluorescence staining

Cells were washed by adding PBS to the apical and basal compartments (transwells) or wells (chamber slide), then fixed for 30 min in 4% PFA (ProSciTech; C005-100) at room temperature (RT). For Zombie Red staining, cells were incubated with a Zombie Red Fixable Viability dye (BioLegend; 423109) according to the manufacturer’s instructions. Briefly,

Zombie Red was added to the apical and basal compartments and cells were incubated for 15 min at RT. Cells were then washed 1X with PBS and fixed for 30 min with 4% PFA in the apical and basal compartments, in the dark. Cells were then blocked and permeabilised for 30 min with 3% BSA (Scientifix; BSASAU) and 0.3% Triton X-100 (Sigma Aldrich; T8787) diluted in PBS +/+ (PBS supplemented with 1□mM CaCl2 and 0.5□mM MgCl2). Cells were stained with primary antibodies against human ICAM-1 (R&D Systems; RDSBBA3), VE-cadherin (R&D Systems; RDSAF938), MUC5AC (Thermo Fisher Scientific; MA512178), acetylated tubulin (Sigma Aldrich; T6793) and nucleocapsid protein (NP) nanobody (NP kindly gifted by Dr. Ariel Isaacs, Watterson Laboratory, School of Chemistry and Molecular Biology, University of Queensland). Primary antibodies were incubated in blocking buffer for 1h at RT. Secondary antibodies were all linked to Alexa fluorophores (all Thermo Fisher Scientific). Cells were stained with secondary antibodies and AlexaFluor 670 Phalloidin (Cytoskeleton, Inc.; CYT-PHDN1-A) in blocking buffer for 1h at RT. Cells were then stained with DAPI (Sigma Aldrich; D9542) in PBS+/+ for 5 min at RT. Cells were washed 3 times between staining steps with PBS+/+ and mounted with ProlongGold (New England Biolabs; 9071S). Z□stack images were acquired on Zeiss LSM 880 Point Scanning confocal microscope using a 40X NA 1.3 water immersion objective. All representative images are maximum image projections generated from Z-stacks. All microscopy imaging was performed at the Institute for Molecular Bioscience Microscopy Facility.

### H&E staining

Undifferentiated (non-airlifted) NHBEs and NHBE/HMVEC-L co-cultures were fixed with 4% PFA overnight at 4°C, dehydrated through graded ethanols, cleared in xylene, and embedded in paraffin. Sections (5 μm) were cut using a microtome and mounted on glass slides. Slides were deparaffinised, rehydrated, and stained with hematoxylin and eosin (H&E) using a standard protocol with Mayers Haematoxylin (Sigma-Aldrich; MHS32) and Eosin Y (Sigma-Aldrich; E4009). Images were acquired on a Zeiss AxioImager 2 upright microscope using a 20X objective.

### Immunohistochemistry

Lungs were harvested and fixed in either 4% paraformaldehyde (PFA) or 10% neutral-buffered formalin for 24h, followed by 70% ethanol dehydration, paraffin embedding, and sectioning. 5 μm thick sections were deparaffinised in xylene and rehydrated through graded ethanol washes. For fibrinogen staining, antigen retrieval was performed in Tris-EDTA buffer (10□mM Tris, pH 9.0, 1□mM EDTA) for 30 min at 95□°C, followed by incubation with proteinase K for 5□min at RT. The lung tissue sections were blocked with 10% BSA in TBS for 1h at RT and then incubated overnight at 4□°C with rabbit anti-fibrinogen/fibrin (Dako; A 0080). Antigen detection was performed using a Zytochem-Plus AP Polymer Kit per the manufacturer’s instructions (Zymed Systems, POLAP-006), and the slides were imaged using an Axio Imager Z2 microscope on a 20x objective (Zeiss).

For ICAM-1 staining, antigen retrieval was performed using Tris-EDTA buffer for 15 min at 95 °C. Lung sections were washed once with MilliQ water, then blocked (20% FBS, 1% BSA in TBST-X) for 1h at RT. Sections were then incubated overnight at 4 °C with primary antibodies against mouse ICAM-1 (BioXCell; BE0020-1) and mouse CD31 (New England Biolabs; 77699S). Sections were then extensively washed with TBS and incubated with secondary antibodies conjugated to Alexa fluorophores (Thermo Fisher Scientific) for 1□h at RT. After washing with TBS, slides were mounted with Prolong Gold (New England Biolabs; 9071S). Z□stack images were acquired on a confocal laser scanning microscope (Zeiss LSM880) using a 20X NA 0.8 objective.

### Isolation of human platelets and platelet adherence assay

Human platelets were isolated from healthy donor whole blood as described previously^55^. Briefly, all centrifugation steps were performed without a brake. Whole blood was centrifuged for 10 min at 330 x g to obtain platelet-rich plasma (PRP). To prevent platelet activation, the following isolation steps were performed in the presence of 200 nM Prostaglandin E1 (PGE1) (Sigma Aldrich; P5515). Two-thirds of the PRP was transferred to a fresh tube and diluted 1:1 with PBS, then centrifuged for 10 min at 240 x g. PRP was transferred to a new tube and centrifuged at 430 x g for 15 min to pellet platelets and remove leukocytes. Platelets were then washed 1X with PBS and labelled with CellTrace Yellow (Thermo Fisher Scientific; C34567), according to the manufacturer’s instructions. Co-culture, triple culture, or HMVEC-L monoculture transwells were inverted into a 6-well cell culture plate, and 1 x 10^6^ platelets were added to the basal side (onto HMVEC-L). Platelets were incubated with HMVEC-L for 30 min at 37°C, 5% CO_2_. Transwells were then re-inverted and washed 1X with PBS to remove non-adherent platelets, before fixation with 4% PFA in the apical and basal compartments for 30 min in the dark.

### RNA extraction and quantitative PCR (qPCR)

Cells were lysed in Buffer RLT plus β-mercaptoethanol, and the RNA was directly processed using an RNeasy Micro Kit (QIAGEN; 74004) with on-column DNase digestion according to the manufacturer’s instructions. RNA concentration was measured using a NanoDrop spectrophotometer (Thermo Fisher Scientific), with equal starting concentrations of RNA for each sample used for reverse transcription. Reverse transcription was performed using Superscript III Reverse Transcriptase (Thermo Fisher Scientific; 18080085) with random nonamer priming. Quantitative polymerase chain reaction (qPCR) was performed using SYBR green reagent (Thermo Fisher Scientific; 4312704) on a QuantStudio 7 Flex Real-Time PCR System (Thermo Fisher Scientific) in 384-well plates. Relative gene expression was determined using the change-in-threshold (2-ΔΔCT) method with hypoxanthine phosphoribosyltransferase 1 (HPRT) as an endogenous control. Primers are as follows: HPRT forward: TCAGGCAGTATAATCCAAAGATGGT, and reverse: AGTCTGGCTTATATCCAACACTTCG; FoxJ1 forward: GATCACGGACAACTTCTGCTACTTC, and reverse: GAGACAGGTTGTGGCGGATT.

### Cytokine analysis

Cytokine levels in cell culture supernatants were measured using the human TNF or IL-1β AlphaLISA Immunoassay kit (Revvity; AL3157C, AL3160C), according to the manufacturer’s instructions, and analysed on a VICTOR Nivo Plate Reader (Revvity).

### Confocal image analysis

Five random regions of interest (ROIs) per sample were acquired for image analysis. Each group included one sample per independent experiment, and the total number of independent experiments is specified in the corresponding figure legends. Raw images were analysed using Fiji (1.54m). For ICAM-1 intensity analysis, max intensity projections were generated from Z-stacks. A mask of the total cell area was created using MaxEntropy thresholding of ICAM-1 staining, and the raw integrated density (RawIntDen) of ICAM-1 within the mask was measured. Values were then corrected for total mask area, resulting in RawIntDen/Area.

To quantify the percentage of the endothelial monolayer area with intercellular gaps, a mask of cell coverage was created using VE-cadherin signal intensity. Any VE-cadherin negative areas were then analysed as particles in Fiji (size=4.00-Infinity, circularity=0.20-1.00). Next, the total number and area of these gaps (measured in mm) per ROI were quantified and used to calculate the total gap area per ROI. HMVEC-L gap quantification is presented as the percentage of gap area relative to total image area.

To quantify the percentage of NP^+^ and Zombie Red^+^ cells, the total number of cells, NP^+,^ and Zombie Red^+^ cells were first quantified using the Spots module of Imaris (Bitplane, version 9). The estimated XY diameter was set to 6 μm and 8 μm for NHBE and HMVEC-L cells, respectively. Background subtraction and quality filter were applied, with the threshold adjusted manually for NHBE cells and automatically for HMVEC-L cells.

For platelet adhesion quantification, immunofluorescent images were analysed using Fiji. Max-intensity projections were generated from Z-stacks, and an ROI encompassing the entire cell area was created using the default intensity threshold for VE-cadherin staining. MaxEntropy thresholding of platelets was performed to remove background before analysing particles within the ROI and the whole image. Particles within the ROI (on endothelial cells) were subtracted from the particles in the entire image to obtain a count for the particles within monolayer gaps. Particles smaller than 2 µm were excluded.

### Immunohistochemistry image analysis

Five random regions of interest (ROIs) per mouse were acquired for image analysis. Raw images were analysed using Fiji (1.54m). For ICAM-1 intensity analysis, max intensity projections were generated from Z-stacks. To measure ICAM-1 intensity on endothelial cells, a mask of the total CD31^+^ endothelial cell area was created using MaxEntropy thresholding of CD31 staining, and the raw integrated density (RawIntDen) of ICAM-1 within the mask was measured. Values were then corrected for total mask area, resulting in RawIntDen/Area.

Fibrinogen staining was analysed in ImageJ by quantifying relative intensity across the entire image and is presented as fibrinogen intensity/mm2.

### Statistical analysis

All statistical analyses were performed using GraphPad Prism version 10 (GraphPad Software Inc., San Diego, California). Data are presented as mean□±□SEM with individual data points indicated and colour-coded per independent experimental replicate. All data were tested for normality using the Shapiro–Wilk test. Specific statistical tests used are detailed in the figure legends. \**p*□<□0.05, \*\**p*□<□0.01, \*\*\**p*□<□0.001 and \*\*\*\**p*□<□0.0001.

## Acknowledgements

This work was supported by National Health and Medical Research Council (NHMRC) of Australia funding, including grants (2010757 to L.I.L, E.J.G, and K.Y.C; 2007979 to K.R.S.). L.I.L. is funded by the Australian Research Council (Future Fellowship FT240100816). M.D. is supported by a Bellbery-Viertel Senior Medical Research Fellowship. M.D.J is supported by a UTS Chancellor’s Research Fellowship. Cell imaging was performed at the Institute for Molecular Bioscience’s Microscopy Facility.

## Author Contributions

K.R.S., K.Y.C, E.J.G, and L.I.L conceived the study and acquired project funding. H.M. performed and analysed *in vitro* co-culture experiments and, together with L.I.L., prepared the final figures. B.H., Y.Z., and A.Y. performed and analysed *in vitro* co-culture experiments. M.D.J. and C.N. performed *in vivo* experiments (K18-ACE2 mice) with input from P.M.H., while S.M.B. and M.D performed *in vivo* experiments with mouse-adapted SARS-CoV-2 (P21 strain) and specific knockout mice. K.R.S, K.R.C., E.J.G., and L.I.L. wrote the manuscript, with input from H.M. E.J.G and L.I.L. supervised the project, and L.I.L. managed collaborations. During the preparation of this work, the authors used Grammarly to improve the manuscript’s readability. After using this tool/service, the authors reviewed and edited the content as needed and take full responsibility for the content of the published article.

## Figures and Figure Legends

**Supplementary Figure 1. Establishment and characterisation of NHBE/HMVEC-L co-cultures. A)** Schematic of NHBE/HMVEC-L co-cultures. **B)** Hematoxylin and eosin staining of undifferentiated NHBEs (top) and fully differentiated (4-week) NHBE/HMVEC-L co-cultures (bottom). Scale bar = 50 mm. **C)** *FOXJ1* mRNA expression was quantified relative to the housekeeping gene *HPRT* in undifferentiated versus 4-week differentiated NHBEs. n = 2 independent experiments. **D)** Immunofluorescent staining of undifferentiated and 4-week differentiated NHBEs for acetylated Tubulin and Muc5AC expression. YZ projection shows a merged image of acetylated tubulin, F-actin, and DAPI, with the apical side at the top and the basolateral side at the bottom. Scale bar = 50 μm. **E)** NHBE/HMVEC-L co-cultures or HMVEC-L monocultures were infected with SARS-CoV-2 (MOI 1). At 72 h post-infection, cells were fixed and stained for the SARS-CoV-2 nucleocapsid protein (NP). Scale bar = 50 µm. Images are representative of n = 4 independent experiments. **F)** The percentage of NP cells was quantified relative to total cells, across 5 ROIs (small dots) per 4 independent experiments (mean = large circles), and analysed by Kruskal-Wallis test with Dunn’s multiple testing correction. **G)** Viral titres (shown as Log_10_ PFU/mL) in supernatants from SARS-CoV-2-infected NHBE/HMVEC-L co-cultures (apical and basal) and HMVEC-L monocultures, at 0 and 72h post-infection, and shows n = 4 independent experiments analysed by unpaired, two-sided Mann-Whitney test. All data are presented as mean ± SEM with individual data points indicated and colour-coded per independent experimental replicate. Asterisks indicate statistical significance: * *p* < 0.05, ** p < 0.01, *** p < 0.001.

**Supplementary Figure 2: HMVEC-L exposed to SARS-CoV-2 maintain functional phenotypes.** (**A-F)** HMVEC-L monocultures were infected with MOI 1 SARS-CoV-2 or mock-infected for 72h and stained as indicated. Scale bars for all images = 50 µm. 5 ROIs per experiment were quantified (small dots) and are colour-coded per experiment. The average of the 5 ROIs is represented by the large dot (colour-coded by experiment), and the data show the mean ± SEM. **A)** Immunofluorescence staining for ICAM-1 (green), F-actin (Phalloidin; grey), and nuclei (DAPI; blue). Images are representative of n = 6 independent experiments. **B)** ICAM-1 intensity between conditions was analysed by an unpaired, two-sided Mann-Whitney test. **C)** Immunofluorescence staining for VE-cadherin (magenta), F-actin (Phalloidin; grey), and nuclei (DAPI; blue). Images are representative of n = 6 independent experiments. **D)** Quantification of gaps in the endothelial monolayer was determined by calculating the percentage of the image area covered by gaps, and analysed by an unpaired, two-sided Mann-Whitney test. **E)** Zombie Red-stained cells (magenta) indicate cells (containing F-actin and nuclei) undergoing cell death. Images are representative of n = 4 independent experiments. **F)** The percentage of Zombie Red positive cells was analysed by an unpaired, two-sided Mann-Whitney test. **G)** Immunofluorescent staining of CellTrace Yellow labelled platelets incubated with HMVEC-L. Images are representative of n = 3 independent experiments. **H)** The number of platelets (CellTrace Yellow positive particles larger than 2 mm) in gaps was quantified and analysed by an unpaired, two-sided Mann-Whitney test. Asterisks indicate statistical significance: * *p* < 0.05, ** p < 0.01, *** p < 0.001.

**Supplementary Figure 3. TNF and IL-1β, but not IL-6, stimulation phenocopies SARS-CoV-2-induced HMVEC-L dysfunction.** HMVEC-L were stimulated with 100 ng/ml IL-6, 100ng/ml IL-6 and 450ng/ml sIL-6R, 100□ng/ TNF or 10ng/ml IL-1β for 24h before fixation. **A)** Immunofluorescence staining for ICAM-1 (green), F-actin (Phalloidin; grey), and nuclei (DAPI; blue). Images are representative of n = 2 independent experiments. **B)** ICAM-1 intensity between conditions was quantified. **C)** Immunofluorescence staining for VE-cadherin (magenta), F-actin (Phalloidin; grey), and nuclei (DAPI; blue). Images are representative of n = 2 independent experiments. **D)** Quantification of gaps in the endothelial monolayer under different conditions was determined by calculating the percentage of the image area covered by gaps. **E)** Zombie Red-stained cells (magenta) indicate cells (containing F-actin and nuclei) undergoing cell death. Images are representative of n = 2 independent experiments. **F)** The percentage of Zombie Red positive cells was quantified. **G)** Immunofluorescent staining of CellTrace Yellow labelled platelets incubated with HMVEC-L. Images are representative of n = 2 independent experiments. **H)** The number of platelets (CellTrace Yellow positive particles larger than 2 mm) in gaps was quantified.

**Supplementary Figure 4. 400 pg/ml TNF stimulation phenocopies SARS-CoV-2-induced HMVEC-L dysfunction.** HMVEC-L were stimulated with 400□pg/ml TNF. **A)** Immunofluorescence staining for ICAM-1 (green), F-actin (Phalloidin; grey), and nuclei (DAPI; blue). Images are representative of n = 2 independent experiments. **B)** ICAM-1 intensity between conditions was quantified. **C)** Immunofluorescence staining for VE-cadherin (magenta), F-actin (Phalloidin; grey), and nuclei (DAPI; blue). Images are representative of n = 2 independent experiments. **D)** Quantification of gaps in the endothelial monolayer under different conditions was determined by calculating the percentage of the image area covered by gaps. **E)** Zombie Red-stained cells (magenta) indicate cells (containing F-actin and nuclei) undergoing cell death. Images are representative of n = 2 independent experiments. **F)** The percentage of Zombie Red positive cells was quantified. **G)** Immunofluorescent staining of CellTrace Yellow-labelled platelets incubated with HMVEC-L. Images are representative of n = 2 independent experiments. **H)** The number of platelets (CellTrace Yellow positive particles larger than 2 mm) in gaps was quantified.

