## Supplementary figures and images for "IL-1β and TNF drive endothelial dysfunction and coagulopathy in acute COVID-19"

### Supplementary Figure 1

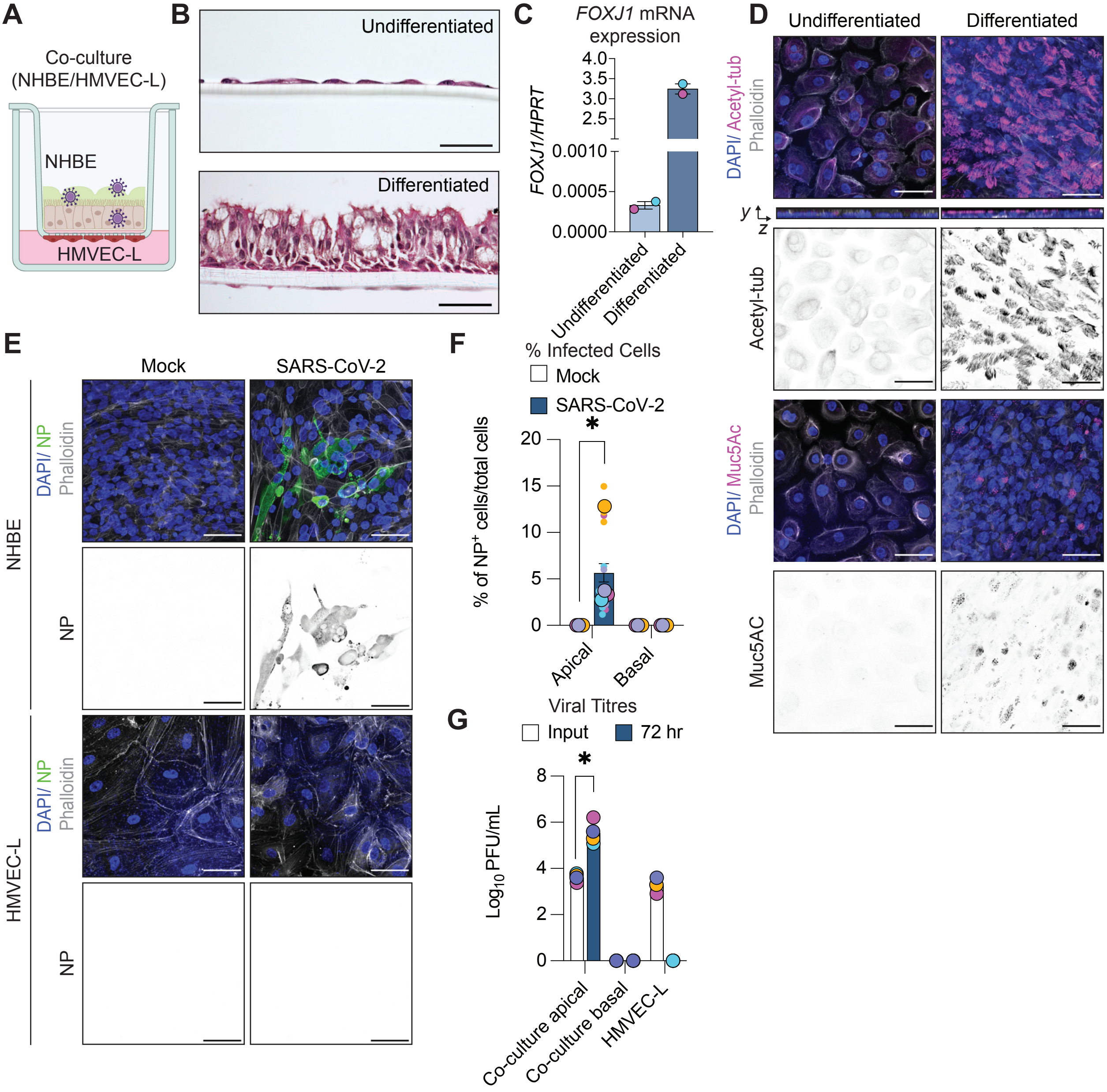

### Supplementary Figure 2

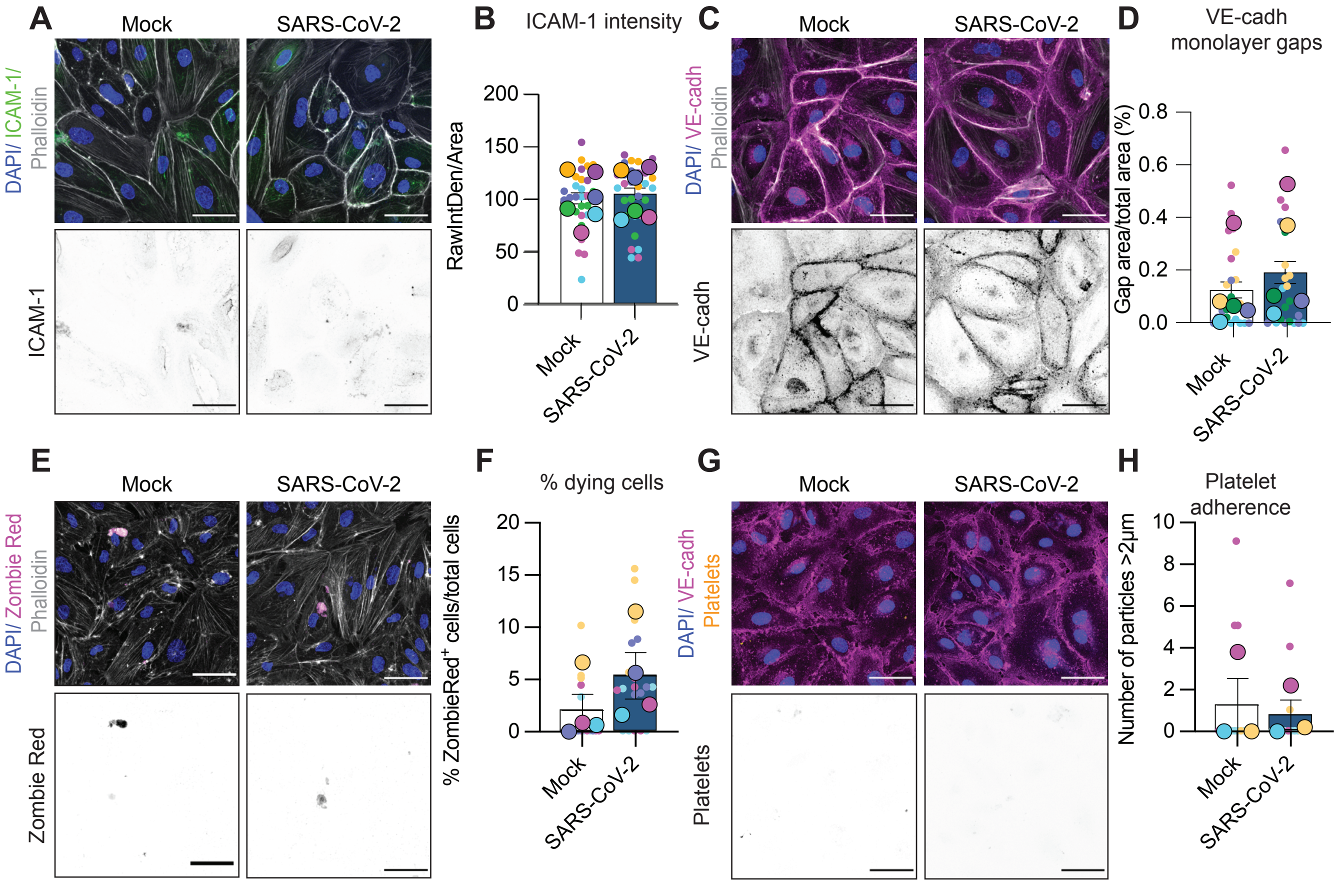

### Supplementary Figure 3

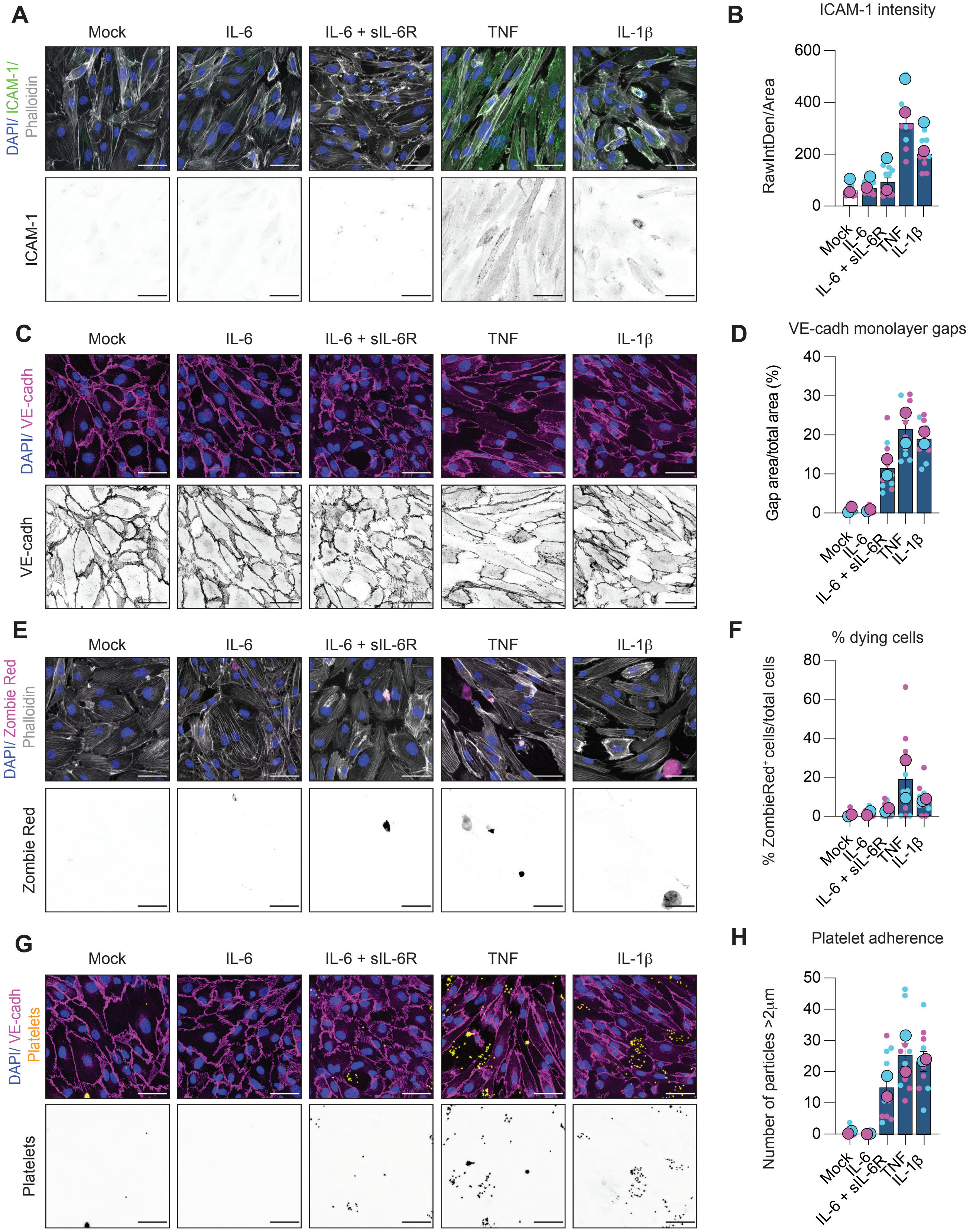

### Supplementary Figure 4

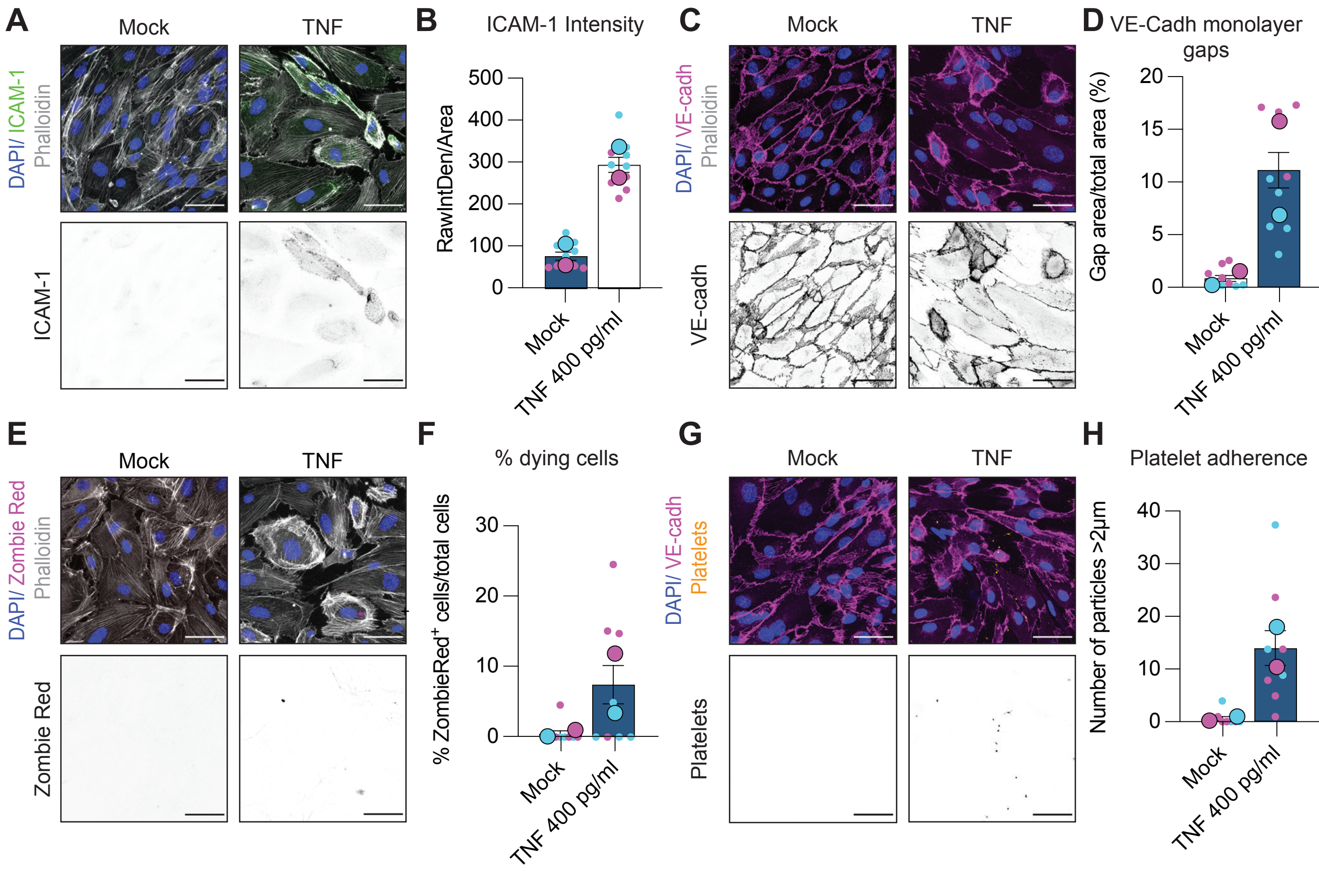
